# A canstatin-derived peptide provides insight into the role of Capillary Morphogenesis Gene 2 in angiogenic regulation and matrix uptake

**DOI:** 10.1101/705459

**Authors:** Jordan G. Finnell, Tsz-Ming Tsang, Lorna Cryan, Samuel Garrard, Sai-Lun Lee, P. Christine Ackroyd, Michael S. Rogers, Kenneth A. Christensen

## Abstract

Capillary Morphogenesis Gene 2 protein (CMG2) is a transmembrane, integrin-like receptor and the primary receptor for the anthrax toxin. In addition to its role as an anthrax toxin receptor, CMG2 has been repeatedly shown to play a role in angiogenic processes. However, the molecular mechanism mediating observed CMG2-related angiogenic effects has not been fully elucidated. Previous studies have found that CMG2 binds type IV collagen (Col-IV), a key component of the vascular basement membrane, as well as other ECM proteins. Currently, no link has been made between these CMG2-ECM interactions and angiogenesis; however, ECM fragments are known to play a role in regulating angiogenesis. Here, we further characterize the CMG2-Col-IV interaction and explore the effect of this interaction on angiogenesis. Using a peptide array, we observed that CMG2 preferentially binds peptide fragments of the NC1 (non-collagenous domain 1) domains of Col-IV. These domains are also known as the fragments arresten (from the α1 chain) and canstatin (from the α2 chain) and have documented antiangiogenic properties. A second peptide array was probed to map a putative binding epitope. A top hit from the initial array, a canstatin-derived peptide, binds to the CMG2 ligand-binding von Willebrand factor A (vWA) domain with sub-micromolar affinity (peptide S16, K_d_ = 400 ± 200 nM). This peptide competes with anthrax protective antigen (PA) for CMG2 binding, and does not bind CMG2 in the presence of EDTA. Together these data suggest that, like PA, S16 interacts with CMG2 at the metal-ion dependent adhesion site (MIDAS) of its vWA domain. We demonstrate that CMG2 specifically mediates endocytic uptake of S16, since CMG2-/- endothelial cells show markedly reduced S16 uptake, without reducing total endocytosis. Furthermore, we show that S16 reduces endothelial migration but not cell proliferation. Taken together, our data demonstrate that a Col IV-derived anti-angiogenic peptide acts via CMG2, suggesting a possible link between CMG2-Col IV interactions and angiogenesis.

## Introduction

CMG2, also known as ANTXR2,^1^ is an integrin-like receptor that was originally identified as a gene that is highly upregulated during endothelial cell tube formation *in vitro.^1^* Like integrins, CMG2 contains an extracellular vWA^1^ domain that coordinates a divalent metal ion in a MIDAS^1^ domain and binds various extracellular matrix proteins. The initial report qualitatively showed CMG2 binding to Col-IV,^1^ fibronectin, and laminin;^*1*^ subsequent reports indicate that CMG2 also binds Col VI.^*2*^ However, the affinity, specific binding sites on each ECM, and the cellular relevance of these interactions have only been superficially examined.

CMG2, like its close homolog TEM8, is most studied its role as an anthrax toxin receptor. These two cellular receptors bind the non-toxic anthrax toxin subunit, protective antigen (PA), and subsequently mediate intracellular delivery of the catalytic toxin subunits.^3, 4^ Experiments that challenged mice lacking full-length CMG2 or TEM8 with *B. anthracis* spores indicate that CMG2 is the major receptor of anthrax toxin^5^ Biophysical characterization also demonstrates that PA has a substantially higher affinity (100 fold) for CMG2 than for TEM8.^6, 7^

While interaction of CMG2 with PA does not illuminate the role of CMG2 in the absence of anthrax toxin, the tight interaction of CMG2 with PA has been used to probe the role of ligand interaction with CMG2 in angiogenesis. Notably, the binding of a mutant PA, (PA-SSSR) to CMG2 substantially inhibits growth-factor induced angiogenesis in the cornea, reduces tumor volume *in vivo*, and blocks endothelial cell migration *ex vivo.^8^* These data indicate that interaction of ligands with CMG2 has a profound effect on angiogenic processes. Further, knockdown of CMG2 in endothelial cells has been reported to inhibit their proliferation and tubule formation,*^9^* and several additional studies have demonstrated the relevance of CMG2 targeting in the inhibition of angiogenesis.*^10–13^*

Substantial data indicate that CMG2 is also important for extracellular matrix (ECM) homeostasis. Mutations in CMG2 that result in loss of CMG2 function are responsible for a severe genetic disorder known as hyaline fibromatosis syndrome, or HFS.^*14–17*^ HFS is characterized by aberrant accumulation of hyaline material under skin and in other organs. Recently, wild-type CMG2 was found to mediate cellular uptake and clearance of type VI collagen (which accumulates in HFS patient nodules), without affecting mRNA levels.*^2^*

There has previously been no functional connection between the role of CMG2 in ECM homeostasis and the role of CMG2 in angiogenic regulation. However, remodeling of the vascular basement membrane (VBM, composed predominantly of Col-IV, laminins, Col-VI, and other glycans) is an essential step in angiogenesis.^*18*^ Hence, it is possible that CMG2 may regulate angiogenesis through VBM/ECM remodeling, which may occur in part via uptake of ECM materials, including protein hydrolysis products. This hypothesis is consistent with the observation that CMG2 endocytoses Col-VI and/or Col-VI fragments, and with buildup of Col-VI in HFS patients that lack functional CMG2.*^2^*

Individual ECM fragments themselves can impact angiogenesis. Specifically, liberated Col-IV C-terminal NC1 domains are potently anti-angiogenic.*^19–21^* These domains appear to act in a negative feedback loop: as angiogenesis proceeds, the VBM is remodeled to allow vessel outgrowth, while at the same time newly-generated Col-IV NC1 fragments engage endothelial receptors to suppress further vessel sprouting.^*22*^ Importantly, Col-IV NC1 domains are essential for both proper triple-helix formation (through NC1 domain trimerization) and ECM network formation (through dimerization of adjacent NC1 trimers). While the 6 distinct Col-IV chains can trimerize in 3 different combinations, the most abundant Col-IV isoform, including within the VBM, is composed of two α1 chains and an α2 chain. The NC1 domains of the α1 and α2 chains are the angiogenesis inhibitors arresten and canstatin, respectively.

Aspects of the mechanism(s) by which arresten and canstatin inhibit angiogenesis have been described, including identification of specific integrin receptors and downstream signaling pathways.*^22–24^* However, the ultimate fate of these Col-IV NC1 domains, including potential receptor-mediated endocytic and degradation pathways, has not been outlined. Given both the established interaction between CMG2 and Col-IV, and the involvement of CMG2 in both ECM homeostasis and angiogenic processes,*^1, 2, 9, 12^* it is possible that peptide fragments of these domains interact with CMG2 and are taken into cells via interaction with CMG2. If so, this interaction could be in part responsible for the anti-angiogenic effects of CMG2 targeting. Hence, we undertook investigations designed to determine whether possible proteolytic peptide fragments of the NC1 domain could interact with cell-surface CMG2 and thereby influence angiogenesis.

Here, we report and characterize a novel interaction between CMG2 and peptide fragments of canstatin. This unanticipated interaction was initially observed via analysis of overlapping peptide arrays of the Col-IV α1 and α2 sequences, which identified peptide sequences in the Col-IV NC1 domain as possible CMG2 binding sites. Peptide array hits with the highest binding intensity were synthesized for further analysis and characterization of their anti-angiogenic effects. Significantly, a canstatin-derived 15-mer peptide (denoted as S16) exhibited both high affinity for CMG2 and potent blocking of endothelial cell migration, suggesting that this small peptide can mimic the anti-angiogenic behavior of full-length NC1 domains. CMG2 mediates endocytosis and perinuclear-lysosomal delivery of this peptide fragment, consistent with a role for CMG2 in ECM/VBM fragment clearance. These findings suggest that CMG2 may play an important role in the regulation of angiogenesis by Col-IV NC1 fragments, and that uptake and degradation of these antiangiogenic VBM fragments could be a functional explanation for the pro-angiogenic behavior of CMG2.

## Materials and Methods

### Protein preparation

Human CMG2 vWA-GST (CMG2-GST), CMG2 vWA R40c and C175A, and PA were expressed and purified as previously described^*25*^. Briefly, CMG2 vWA (40-217) was expressed with R40C and C175A mutations (R40C provides an exposed site for maleimide labeling; and C175A removes a buried cysteine to ensure a 1:1 labeling ratio) from a pGEX-4T1 vector. Human CMG2-GST was expressed in BL21 T7 Express *E. coli* (New England Biolabs) with 0.5 mM IPTG induction in a 5L bioreactor (Sartorius). Cells were lysed via French press and sonication, and CMG2-GST was affinity purified with Glutathione Superflow Agarose (Thermo Fisher). Glutathione was removed and CMG2 exchanged into HBS-T with 50% glycerol using Sephadex G50 (GE Life Sciences) on an Äkta Start chromatography system.

His tag-mCitrine-TEM8 vWA was expressed and purified as described previously.^*26*^ Briefly, His tag-mCitrine-TEM8, from a pQE30 vector, was expressed in BL21 T7 Express *E, coli* (New England Biolabs) with 1mM IPTG induction in a shake flask with 1L LB supplemented with 10mM Glucose. Cells were lysed in 10mM imidazole PBS via French press to obtain clear lysate, then purified with a Nickel column. Column was washed with 8 column volume of 20mM imidazole PBS, then eluted with 4 column volume of 250mM imidazole PBS. Eluted protein was concentrated by a 30kDa centrifugal filter unit (Millipore Sigma Cat# ufc903024), and stored in 50% glycerol PBS at −20°C.

PA-SSSR and PA-E733C were expressed from pET-22b (RRID: Addgene 11079) into the periplasm and purified from the periplasmic lysate via anion exchange chromatography (Q-sepharose, GE Life Sciences Cat: 25236), using 20 mM Tris-HCl pH 8.0 with 20 mM NaCl (Buffer A) and Buffer A + 1 M NaCl. Endotoxin was removed by passing twice through poly-lysine coated cellulose beads (Thermo Fisher Cat: 88275) followed by an endotoxin test using the gel clot method and appropriate dilutions.

CMG2-GST (AlexaFluor 488 or biotin) and PA-E733C (AlexaFluor 546) were conjugated on a single reactive cysteine with maleimide-labels (Thermo Fisher) after GST was cleaved by using thrombin. Label:protein ratio was 5:1 in the reaction mixture. Conjugated protein was then purified from free label by size exclusion (Sephadex G50). All protein stocks were stored at −80°C in 50% glycerol.

Peptide S16 (PAIAIAVHSQDVSIP) was synthesized (Genscript and Biomatik). Peptide U12 (VSIGYLLVKHSQTDQ) and scrambled S16 peptide (SPIAVDVQSAPIHAI) were synthesized (Genscript). S16 conjugated at the N-terminus to HiLite-488 was also synthesized (Anaspec).

### Membrane-based peptide array with 10 residue sliding window

An overlapping peptide array (15 residue peptides, with a 5-residue overlap) containing the Col-IV α1 (Uniprot P02462) and α2 (Uniprot P08572) sequences was created by direct synthesis of peptides onto an amino-PEG cellulose membrane (ABIMED peptide array; Koch Institute Biopolymers and Proteomics Facility). The membrane was blocked in TBST with 1% milk, then probed with 250 nM CMG2-biotin. Bound CMG2-biotin was detected by avidin-HRP. Spot intensities were visually scored from 0 (no observed binding) to 5 (max binding). Hits were quantified and compared by calculating both a hit ratio and weighted hit ratio for each domain. Hit ratio was calculated by dividing the total number of hits by the total number of peptides in the array for that domain; weighted hit ratio divided the sum of the intensity scores of all the peptide hits in each domain by the total number of peptides in the array for that domain.

### PEPperPRINT peptide micro-array with 2-residue sliding window

A peptide array containing the Col-IV α1 and α2 NC1 domains (15 residue peptides with a 13-residue overlap) was printed by PEPperPRINT and probed according to manufacturer recommendations with 2 µM CMG2-GST then anti-GST-DyLight 800 4X PEG (RRID: AB_2537633). The raw TIFF image was analyzed by PepSlide^®^ Analyzer (Sicasys).

From the quantified data and visual inspection, peptide hits flanked by adjacent hits are considered a potential binding surface. Hits with irregularly shaped noise or background were eliminated from further analysis. Epitope mapping molecular graphics were made in the PyMOL Molecular Graphics System, Version 2.0 Schrödinger, LLC.*^27^*

GibbsCluster-2.0*^28, 29^* was used to perform clustering analysis on array hits. Parameters were systematically varied in order to evaluate the effect of motif length and insertions and deletions on the quality of the alignment. A visual representation of these analyses was generated by Seq2Logo.*^30^*

### CMG2-PA FRET assay

Interaction between CMG2 and peptides were measured using an *in vitro* FRET assay to detect competition with PA, as described previously.^*13*^ Briefly, 10 nM each of CMG2-488 and PA-546 were incubated in the presence of peptide of varying concentration, with DMSO as vehicle control. FRET ratio = I_485/590_/ I_548/528_. FRET ratio with 10 mM EDTA was set to 0.

### TEM8-PA FRET assay

Interaction between TEM8 and peptide S16 were measured using *in vitro* FRET assay to detect competition with PA, as described previously.*^26^* Briefly, 250nM mCitrine-TEM8 and 325nM PA-546 were incubated in the presence of peptide of varying concentrations, with DMSO as vehicle control. FRET ratio = I_485/590_/ I_548/528_.

### Bio-layer interferometry

Bio-layer interferometry was used to characterize the interaction of peptides with CMG2. Assay buffer was 50 mM HEPES, 150 mM NaCl, pH 7.2, 0.1% Tween-20, 1 mg/mL BSA, 2 mM CaCl_2_, 1 mM MgCl_2_, 0.02% NaN_3_. First, streptavidin biosensors (ForteBio) were coated with 5-10 µg/mL CMG2-GST-biotin overnight at 4°C. Eight sensors were loaded and run in parallel (6 sensors for S16 binding, 2 for reference controls). Binding assays were performed the following day, using an Octet RED96 biolayer interferometer (ForteBio) and the Octet 8.2 Data Acquisition software. Assays were performed at 30°C and 1000 rpm shaking. The CMG2-GST-biotin loaded sensors were equilibrated in assay buffer (1200 sec) followed by an association step with a serial dilution (1-90 µM) of peptide (300-1200 sec), and a dissociation in assay buffer (600-1800 sec). Binding data for S16 concentrations below 1 µM were not obtainable, due to the low signal/noise ratio at sub-micromolar concentrations. Data was processed and analyzed in the Octet Data Analysis 8.2 software. Processed data was fit to a 1:1 binding model to obtain kinetic and thermodynamic parameters. Residuals were examined to assess quality of fit and no systematic deviation was observed.

### Cell lines and culturing technique

EOMA (CRL-2586, RRID:CVCL_3507) is a murine hemangioma endothelial cell line.*^31^* EA.hy926 (CRL-2922, RRID:CVCL_3901) cells are the result of a fusion of human umbilical vein cells with lung carcinoma cells^*32*^. Both cell lines were cultured in 10% FBS + DMEM and incubated at 37°C in a humidified environment with 5% CO_2_ until ready for passaging.

### Endothelial cell binding assays by flow cytometry

Cells were cultured in well-plates to 30-70% confluence. Cells were incubated in 10% FBS DMEM and treated with 200 nM PA-SSSR when indicated. Cells were co-treated with S16-Alexa Fluor 488 conjugate (2 µM), transferrin-Alexa Fluor 633 conjugate (10 µg/mL ThermoFisher, Cat: T23362), or Dextran 10,000 MW Texas Red (100µM ThermoFisher, Cat: D1863), and incubated at 37°C for indicated times. When indicated, a well plate with staining media supplemented with 20mM HEPES was left on ice for the indicated time. After incubation, cells were washed, trypsinized, and spun down, then resuspended in cytometry buffer (PBS, 1% BSA, 5 mM glucose) and analyzed in an Attune Acoustic Focusing Cytometer, equipped with 488nm and 633nm lasers. Flow experiments that involved with Texas Red conjugated protein were performed with the Beckman Coulter CytoFLEX cytometer that equipped with 405nm, 488nm, 561nm, and 638nm lasers. Cytometry data was analyzed in FlowJo.

### Confocal microscopy to track ligand endocytosis

Cell preparation and staining for confocal microscopy was as described above for flow cytometry, except that cells were seeded in appropriate glass bottom culture dishes. Staining with S16-488 (2 µM) and transferrin-633 (10 µg/mL) was performed at 37°C for 1-3 h, washed three times with serum-free media, then incubated with a low fluorescence media supplemented with ProLong live-cell antifade reagent (ThermoFisher Cat: P36975) at 37°C for at least 1 hour. Live-cell images were acquired on a Leica DMi8 confocal microscope, using 488nm and 633nm lasers. Images were acquired with the Leica Application Suite X (LAS X), deconvoluted in Huygens Essential, and further analyzed in LAS X to examine colocalization.

### Wound-scratch migration assay

EOMA cells were seeded in 96 well plate and cultured in 10% FBS DMEM. Cells were incubated at 37°C in a humidified environment with 5% CO_2_ until completely confluent. A 200 µL pipette tip was used to scratch a wound in the monolayer, followed by three washes with PBS. Cells were then treated with 10% FBS DMEM with or without S16, and serum-free DMEM as negative control. Images were acquired every four hours, and the change in wound area was quantified using ImageJ.

### Cell proliferation assay

15,000 EOMA cells were seeded into each well in a 96 well plate and incubated for 1 h to attach. After cell attachment, media with treatments were added. Ethanol-fixed cells were used as negative control. After a 24 h incubation, 20 µL CellTiter-Blue Reagent (Promega Cat: G8080) was added to each well for 4 h. Fluorescence signal (Ex: 560nm / Em: 590nm) was measured using a BioTek Synergy H2 plate reader. All readings were normalized to the non-treated control.

### CellASIC migration assay

The assay protocol followed the CellASIC ONIX M04G-02 Microfluidic Gradient Plate User Guide. All media put into the plate (except the cell suspension) was filtered through a 0.2 µm syringe filter. EA.hy926 cells (3 × 10^6^ cells/mL) were loaded in and incubated overnight with DMEM + 10% FBS under flow at 37°C. Assays were then performed with a stable gradient of DMEM + 0 – 10% FBS with or without peptide treatment. Brightfield images at 10x magnification were taken every 10 min over 12 h on an Olympus IX73 microscope and the ORCA-Flash 4.0 camera (Hamamatsu). Individual cells were tracked with ImageJ (RRID: SCR003070) manual tracking plugin. Data was transferred to the ibidi Chemotaxis and Migration Tools 2.0 to export endpoint ‘y’ displacement and accumulated displacement. P-values were calculated using Student’s *t*-test and error bars are standard error of the mean (n=40).

## Results and Discussion

### CMG2 – Col-IV interaction occurs preferentially within the anti-angiogenic NC1 domains of Col-IV determined by peptide array

CMG2 has emerged as an important regulator of angiogene*sis,^8, 9, 12^* but there is still little insight into its mechanism. Given reports that CMG2 binds matrix proteins, including Col-IV^1^, and that the Col-IV fragments have anti-angiogenic properties,*^19–21^* we investigated the interaction of the CMG2 vWA domain with Col-IV peptide sequences to shed light on the possible relevance of this interaction for angiogenic regulation. To identify peptide sequences on Col-IV that can interact with CMG2, we constructed a scouting peptide array in which the linear sequences of human Col-IV α1 and α2 chains were directly synthesized onto a nitrocellulose membrane. The array consisted of 15-mer peptide units, with a 10-residue shift between subsequent peptides, and scanned the entire length of the α1 and α2 chains of Col-IV. The array was probed for CMG2 interaction with 250 nM CMG2-biotin and read out with avidin-HRP; spots representing binding of CMG2 to individual peptide sequences were scored with intensities assigned from 0 (min) to 5 (max) (Fig 1a-b, Supplementary Figure 1). When these intensity values were plotted against the complete sequential protein sequence in the Col-IV α1 and α2 chains, we could visualize relative binding in different sections of the sequence (Figure 1a-b). For each chain, we observed sparse and weak CMG2 binding throughout the Col-IV 7S and triple-helical regions, but heavily concentrated binding within the Col-IV NC1 domains (Fig 1, Table 1). We quantified this binding by calculating both a hit ratio and weighted hit ratio for each domain. Hit ratio was calculated by dividing the total number of hits by the total number of peptides in the array for that domain; weighted hit ratio accounted for the relative intensity of peptide hits. When the peptide array binding data were evaluated this way, we observed that arresten and canstatin exhibited much higher hit ratios and weighted hit ratios than their corresponding triple-helical or 7S domains (Table 1). Specifically, hit ratios and weighted hit ratios for arrestin were 14-fold and 3-fold higher, respectively, than the triple-helical domain and 25-fold and 6-fold, respectively, higher than the 7S domain. These observations indicated that CMG2 preferentially binds the NC1 domain-derived sequences on the peptide array, and suggest that CMG2 may bind to one or more sites on the Col-IV NC1 domain.

**Table 1.** Binding profile quantification for CMG2 interaction with Col-IV-derived peptides. Hit ratio was calculated as # hits (intensity > 0) divided by total number of peptides in domain. Weighted hit ratio was calculated as total measured intensity of domain divided by total number of peptides in domain.

|  | <u># Hits</u><br><u>(Total #</u><br><u>Peptides)</u> | <u>Hit Ratio</u> | <u>Weighted</u><br><u>Hit Ratio</u> | <u>Mean Spot</u><br><u>Intensity±</u><br><u>SEM ‡</u> | <u># Hits (Total</u><br><u># Peptides)</u> | <u>Hit Ratio</u> | <u>Weighted</u><br><u>Hit Ratio</u> | <u>Mean Spot</u><br><u>Intensity±</u><br><u>SEM ‡‡</u> |
| --- | --- | --- | --- | --- | --- | --- | --- | --- |
| 7S | 0 (14) | 0 | 0 | 0 ± 0 | 1 (16) | 0.063 | 1.56 | 0.3 ± 0.3 |
| Triple-helical | 5 (127) | 0.039 | 0.551 | 0.05 ± 0.03 | 16 (129) | 0.124 | 2.64 | 0.26 ± 0.07 |
| NC1 | 13 (23) | 0.565 | 13.9 | 1.5 ± 0.3 | 10 (24) | 0.417 | 15.0 | 1.5 ± 0.4 |

‡ Tukeys post-hoc – 7S vs Triple-helical: p = 0.9394, NC1 vs 7S: p < 0.0001, NC1 vs Triple-helical: p < 0.0001
‡‡ Tukeys post-hoc – 7S vs Triple-helical: p = 0.9827, NC1 vs 7S: p = 0.0009, NC1 vs Triple-helical: p < 0.0001

**Figure 1.**
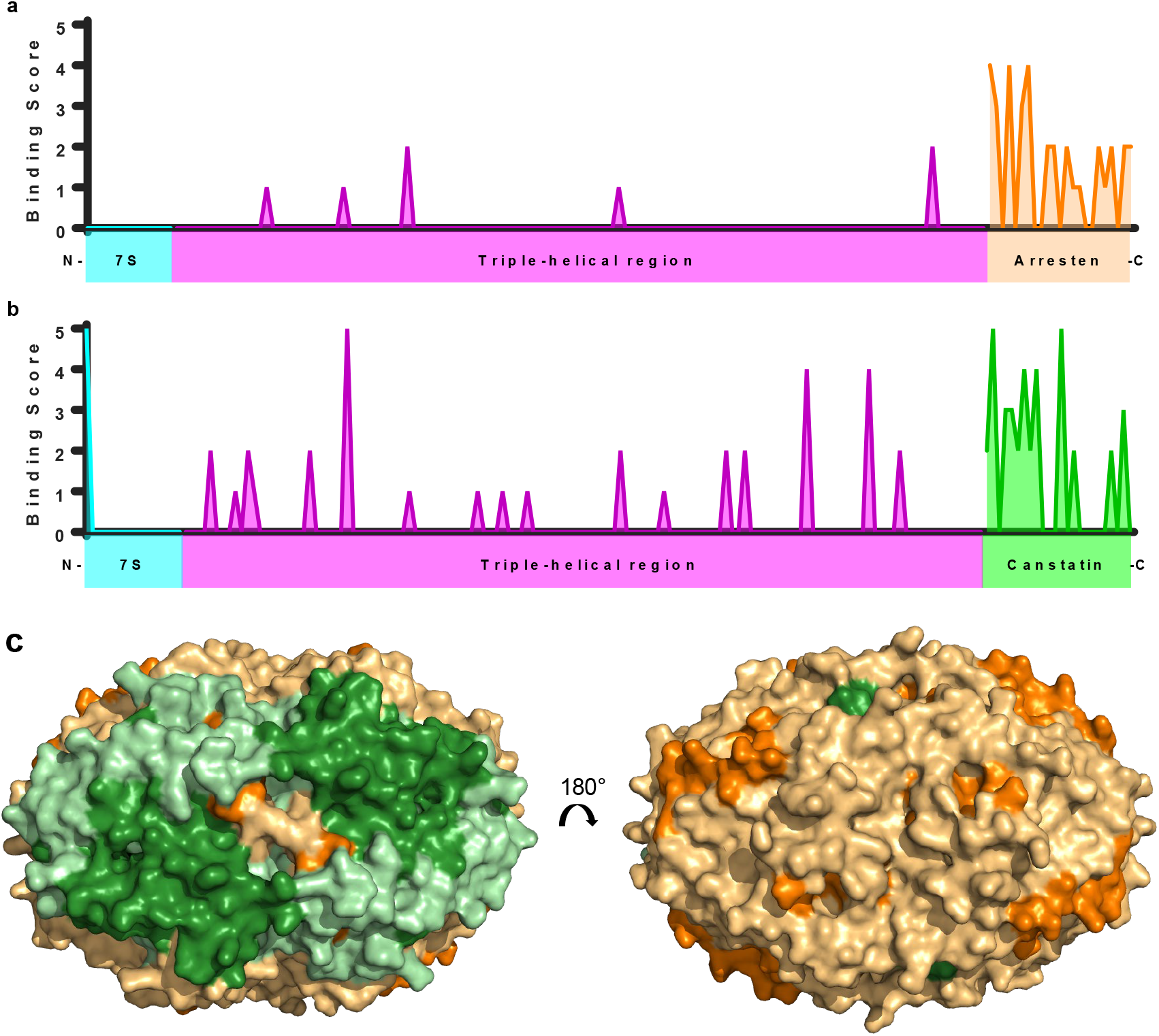
Peptide arrays identify CMG2 as a receptor of Col-IV NC1 domains. (a-b) A peptide array of 15-mer peptides with a 10-residue sliding window, covering the sequences of Col4A1 (a) and Col4A2 (b). This array was probed with 250 nM CMG2-biotin and read out using avidin-HRP. Spot intensities were scored from 0 (no observed binding) to 5 (maximal binding). Domain topology is shown below. There is a greater proportion of CMG2-binding peptides within the c-terminal NC1 domains than the collagenous and 7S domains. (c) PEPperPRINT peptide array with a 2-residue sliding window identifies a putative CMG2-binding surface. NC1 hexamer shown (PDB 1LI1) consists of two trimers, each composed of two arresten and one canstatin. Arresten is shown in orange; canstatin in green. Binding sequences are shown in darker shades. Left shows canstatin-exposed face, demonstrating a large, uniform epitope for CMG2 to canstatin. Right displays arresten-exposed face, where CMG2-binding peptides are present, but are more sparse.

We sought to further characterize the CMG2-NC1 interaction by mapping possible binding site(s) onto the NC1 domain. While the initial peptide array strongly indicated that CMG2 binding is enriched in canstatin and arresten versus the rest of the Col-IV chains, the large sliding window for different peptides in this array made the individual CMG2-binding residues within each peptide hit difficult to identify. To identify the individual peptides in the NC1 domain that could be binding CMG2, we synthesized an additional peptide array on a glass chip, with the NC1 domains arrayed as 15-mer peptides in a 2-residue sliding window (Supplementary Figure 2). The goal was to use hits from the overlapping peptides confirm the identity and sequence of CMG2-binding peptide regions in the NC1 domain. We defined a putative peptide hit as 3 or more consecutive peptides that showing binding to CMG2, and mapped identified CMG2-binding sequences onto a previously solved structure of the Col-IV NC1 domain hexamer (Fig 1c, PDB: 1LI1)^*33*^. Notably, both canstatin and arresten fragment peptides identified as putative CMG2 interactors are localized on the accessible surface of the NC1 domain (Figure 1c, left), and therefore could be recognized by CMG2 for interaction in whole NC1 domain. Peptide fragments from arresten were also identified on the solvent-exposed surface (Fig 1c, right) although the CMG2-binding surface of arresten was sparser than for canstatin, consistent with its lower hit rate. Importantly, identification of individual CMG2 binding peptide fragments of arresten and canstatin, combined with their localization on the solvent accessible surface of the NC1 domain, suggests the possibility that the antiangiogenic activity of arresten and canstatin could be regulated, in part, by interaction with CMG2.

**Figure 2.**
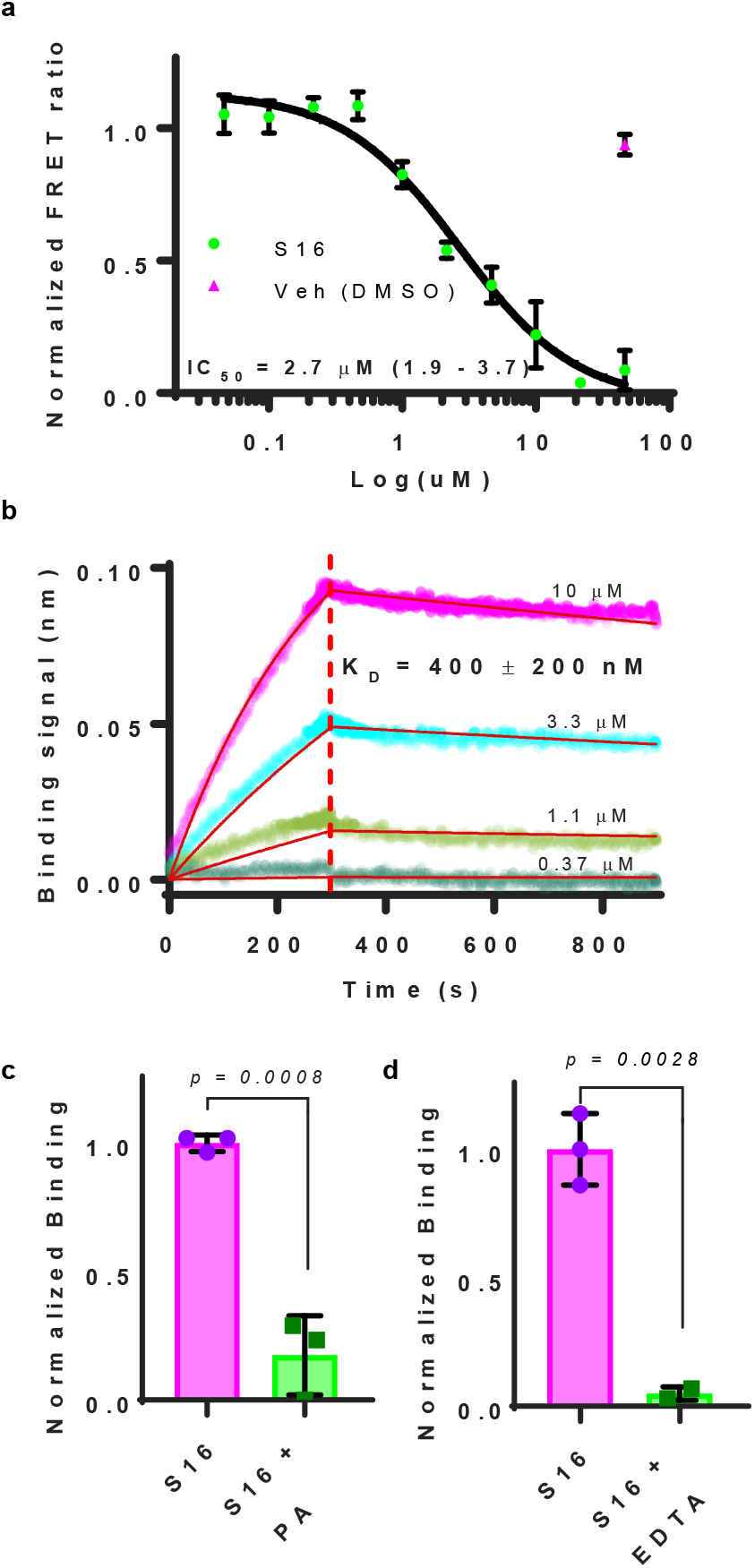
Peptide S16 binds to the CMG2 MIDAS with sub-micromolar affinity. (a) CMG2-488/PA-546 FRET inhibition assay confirms the specificity of S16 binding to CMG2. IC_50_ shown with 95% CI. (b) Representative experiment of S16 (concentration indicated) binding to CMG2-GST-biotin loaded BLI sensors. K_d_ shown is mean ± std (3 independent experiments). (c-d) Peptide S16 interacts with the MIDAS domain of CMG2. (c) CMG2-GST-biotin loaded BLI sensors equilibrated in buffer with or without 500 nM PA, followed by incubation in 10 µM S16 (n=3 per condition). (d) CMG2-GST-biotin loaded BLI sensors were equilibrated either in buffer with 2 mM CaCl_2_ and 1 mM MgCl_2_, or 10 mM EDTA, followed by incubation with 10 µM S16 (n=3 per condition). Error bars are standard deviation of replicates.

CMG2 has high structural homology to many integrins,*^6^* whose ligands commonly have an established RGD motif.*^34^* Hence, we evaluated the 2-residue sliding window peptide array data to identify a possible consistent CMG2-interacting motif. Alignment of all hits from this array produced a consistent binding element distinct from the RGD motif of integrins that contained an aliphatic residue at the n^th^ position, followed by an acidic residue at the (n+2)^th^ position (Supplementary Figure 3). The presence of acidic residues in this motif suggests that, like PA, CMG2 peptide ligands could participate in binding the divalent cation in the CMG2 MIDAS domain.*^35^*

### Canstatin-derived peptide S16 binds with high affinity to CMG2 via the MIDAS

To further understand the relevance of CMG2 interaction with NC1-derived peptides, we subjected the top NC1 hits from the initial peptide array for further evaluation. Peptides from the NC1 region showing the maximum binding score of 5 were selected and characterized using two *in vitro* assays: 1) a PA-CMG2 competition FRET assay,*^13^* and 2) a bio-layer interferometry (BLI) kinetic binding assay. Of the peptides tested, two showed activity in both assays. These peptides are both from canstatin and are denoted here as S16 and U12. Both U12 and S16 bind CMG2 and compete with PA for binding to CMG2 (Figure 2, Supplementary Figure 4). Peptide U12 showed relatively modest affinity for CMG2, both by BLI (K_d_ = 80 µM) and the FRET competition assay with PA-AF546 conjugate (IC_50_ = 35 µM; Supplementary Figure 4). Peptide S16 bound CMG2 with much higher affinity. Analysis by BLI revealed that S16 binds CMG2 with sub-micromolar affinity: K_d_ = 0.4 ± 0.2 μM (Fig 2b, mean ± SD, n = 3 independent experiments). This interaction was characterized by a relatively slow on-rate (≈ 2 × 10^2^ M^−1^s^−1^) and off-rate (≈ 1 × 10^−4^ s^−1^). Further discussion on the observed kinetics is provided in Supplementary Note 1. Binding of CMG2 by S16 was corroborated by FRET competition with PA, in which the S16 IC_50_ was 2.7 μM (Fig 2a, 95% CI: 1.9-3.7), which corresponds to a K_d_ of 100 ± 55 nM, in rough agreement with that measured by BLI.

To confirm the specificity of S16 for CMG2, we synthesized a scrambled S16 peptide comprised of the same residues as S16 but in random order, and probed for interaction of this scrambled peptide with CMG2 via BLI. Unlike S16, the scrambled version of S16 peptide showed minimal binding of CMG2, even at concentrations up to 30 μM (Supplementary Figure 5). Additionally, the scrambled S16 binding data fit poorly to all available binding models tested (1:1, 1:2, and 2:1 bivalent models). These studies are consistent with a sequence-specific interaction between S16 and CMG2.

The ability of S16 to compete with PA for CMG2 binding, as identified in the FRET competition assay, suggests that S16 interacts with the ligand binding surface of the CMG2 vWA domain at the MIDAS, as does PA. BLI experiments were used to verify that S16 binds in competition with PA. Specifically, when CMG2-loaded biosensors were treated with PA prior to exposure to S16, CMG2 binding to S16 was reduced (Fig 2c). As further support of MIDAS interaction, EDTA prevents S16 binding to CMG2 (Fig 2d), confirming that the interaction is metal ion (Ca^2+^ or Mg^2+^) dependent. Indeed, S16 contains an aspartate (preceded by several aliphatic residues) that could participate in metal coordination.

Since CMG2 shares high sequence and structural homology with its competing ANTXR, TEM8*^3, 6^* and both bind PA (albeit with dramatically differing affinities), we asked whether S16 could also bind TEM8. FRET showed that S16 is unable to compete with PA for TEM8 binding, indicating that S16 is not recognized or bound by TEM8 (Supplementary Figure 6). The conclusion that this Col-IV peptide fragment interacts with CMG2 rather than TEM8 is perhaps not surprising; while CMG2 has been reported to bind Col-IV,^*1*^ there is no published evidence for interaction of TEM8 with Col-IV.*^36, 37^*

As described above, multiple independent assays demonstrate specific interaction of peptide S16 with the CMG2 vWA domain. Together, data from these assays demonstrate that this canstatin-derived peptide interacts with the CMG2 MIDAS in a manner competitive with anthrax toxin PA, and with an affinity of ~0.4 μM. Considering the established role of CMG2 in angiogenesis and the anti-angiogenic activity of anthrax toxin PA^*8*^ and the Col-IV NC1 canstatin^*19*^, we asked whether this small fragment of canstatin (S16) can inhibit angiogenic processes in endothelial cells.

### Peptide S16 inhibits endothelial cell migration, but not proliferation

Proliferation and migration of endothelial cells are essential to angiogenesis. We examined the impact of S16 on both proliferation and migration of EOMA cells. Previous work has shown that binding of individual ligands to CMG2, including PA and small molecules, inhibits migration but not proliferation of microvascular endothelial cells.*^8, 10, 38^* Like PA, S16 did not impact endothelial cell proliferation in EOMA cells (Fig 3a). However, in a standard wound scratch assay with EOMA cells, S16 exhibited strong inhibition of endothelial cell migration compared with the untreated, full-serum control (Fig 3b). Indeed, migration of S16-treated EOMA cells was statistically indistinguishable from migration of EOMA cells without serum, indicating that S16 potently reduces cell migration in this cell type. S16 competes with PA for CMG2 binding, and PA is known to inhibit cell migration and other angiogenic processes.^*8*^ Hence, the observation that S16 also inhibits migration suggests that S16 exerts its angiogenic impact via interaction with CMG2.

**Figure 3.**
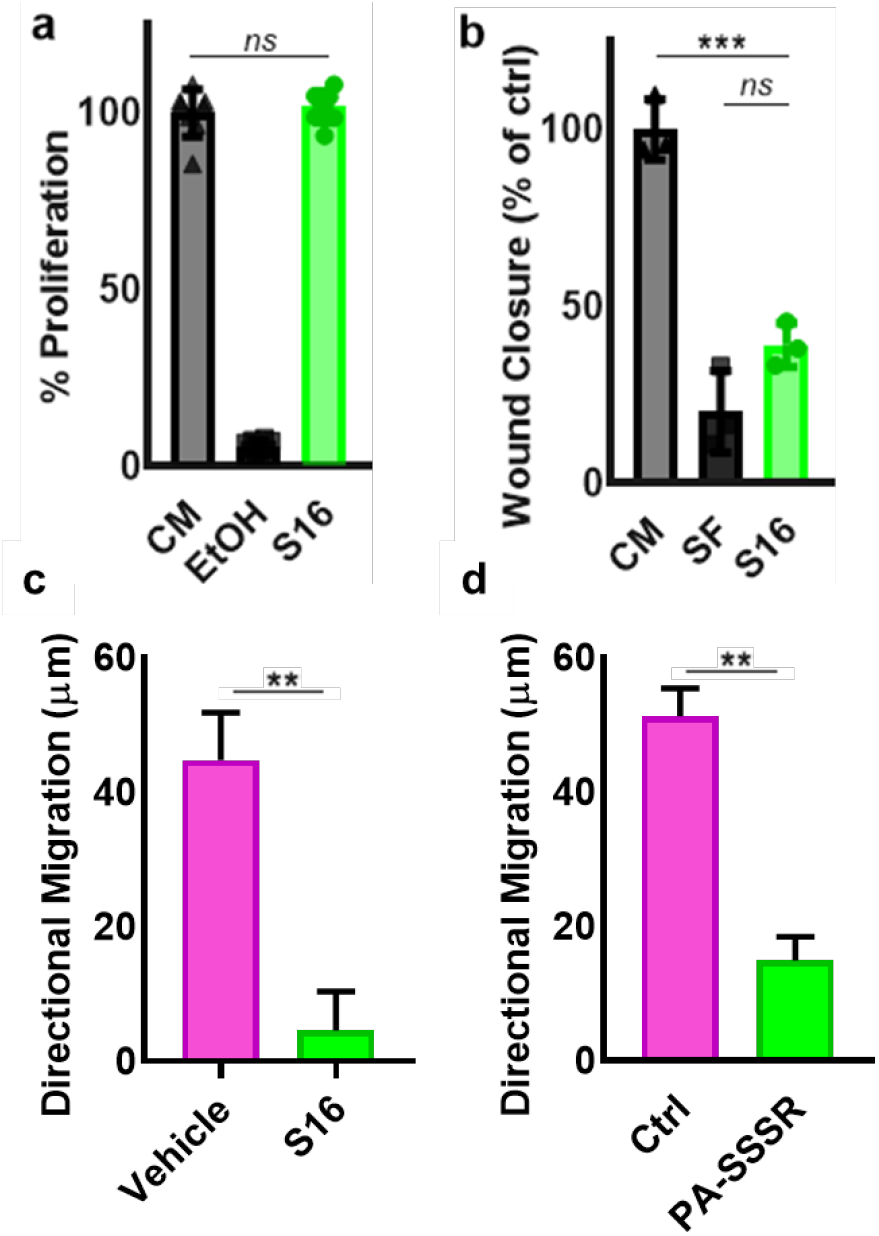
Peptide S16 abolishes migration of endothelial cells, with no effect on proliferation. (a) Proliferation of EOMA cells was monitored. Treatment with S16 resulted in no change in proliferation relative to the untreated control. In contrast, cells fixed in ethanol showed a complete loss of proliferation. Error bars are SD of n=7 replicates from two independent experiments. (b) Wound scratch migration assay. S16 results in 60% reduction in wound closure and is not statistically distinguishable from the serum-free (negative) control. Error bars are SD of n=3 replicates. (c-d) Serum gradient Migration assay with EA.hy926 cells. Data show that cells migrating towards serum in the vehicle control, but a significant loss in migration towards serum with S16 treatment (c) or 200pM PA-SSSR (d). Error bars are SEM of n=40 cells for each condition. **, p<0.01, ***, p < 0.001; ****, p < 0.0001; ns, not significant.

To further characterize the anti-migratory effect of S16 on endothelial cells, we performed additional assays in a microfluidic platform that allows for tracking of individual cell migration within a serum gradient. Real-time tracking of human EA.hy926 cells demonstrated strong migration in the absence of S16 (Fig 3c). In contrast, when S16 was present in the cell medium, migration toward serum was reduced by ~90% (Fig 3c); migration was similarly inhibited in medium containing 200pM PA-SSSR (Figure 3d). This is consistent with the observed lack of mouse cell migration in the wound scratch assay. To confirm that the loss of Ea.hy926 cell migration results specifically from S16 interaction with CMG2, we repeated the gradient migration assay using the same scrambled peptide generated as a CMG2-binding control and found no difference in migration between treated cells and vehicle control (Supplementary Figure 7). This demonstrates that the anti-migratory effect depends specifically on the amino acid sequence of S16, consistent with a requirement for CMG2 binding in the observed anti-migratory effect. Since S16 does not bind TEM8, the anti-migratory effect observed in these studies depends on interaction of S16 with CMG2, rather than with TEM8, which is also present in these cells. Taken together, these data indicate that S16 inhibits endothelial cell migration in a CMG2-dependent manner.

### CMG2 mediates endocytosis of S16 in endothelial cells

Having determined that S16 binds to CMG2 with high specificity and affinity (Fig. 2) and that S16 inhibits endothelial cell migration (Fig. 3), we sought to confirm that, like other CMG2 ligands, S16 is also internalized. We quantified internalized S16 via flow cytometry analysis of Ea.hy926 cells treated with a fluorescently conjugated S16 construct (S16-488), and compared the S16 signal to that of Transferrin-633 (Tf-633), a commonly used endocytic marker. We first compared S16-488 and Tf-633 fluorescence in EA.hy926 cells at 37°C to that of cells treated on ice, a condition that inhibits endocytic pathways including both receptor-mediated uptake and (nonspecific) fluid-phase pinocytosis.*^39^* The robust S16-488 and Tf-633 flow cytometry signal exhibited at 37°C was completely eliminated by incubation on ice (Fig 4a). Hence, S16, like transferrin, is endocytosed by these endothelial cells.

**Figure 4:**
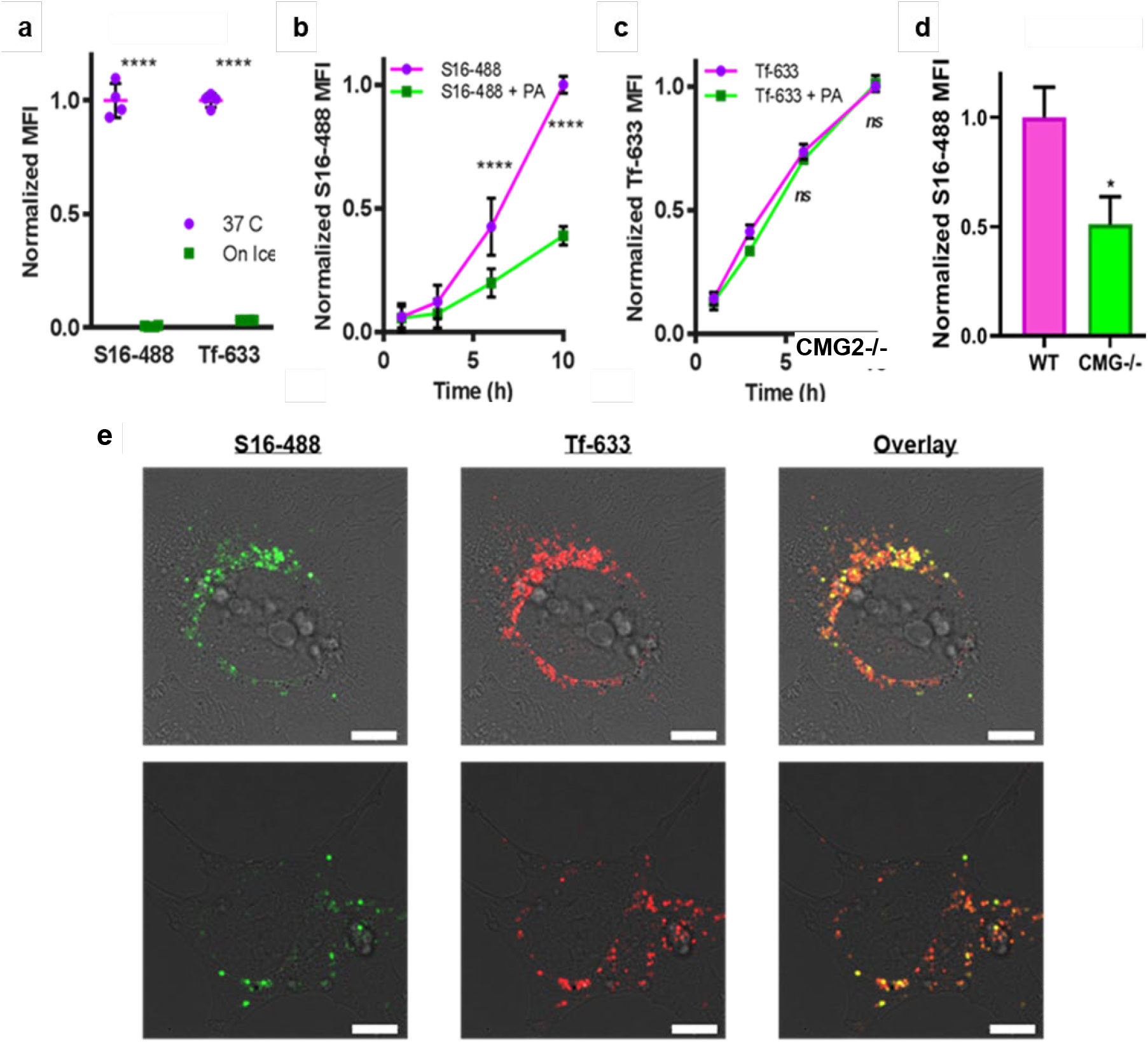
CMG2 is the relevant cell surface receptor for S16 and mediates S16 internalization. (a) Cold treatment abolishes the signal of both S16-488 and transferrin-633, indicating that, as transferrin, S16 is being internalized. Conditioned EA.hy926 cells were incubated with 2 µM S16-488 and 10 ug/mL transferrin-633 in DMEM (with 20 mM HEPES and 1% FBS) for 6 hrs, either at 37C or on ice. Cells were harvested and analyzed by flow cytometry. Plot shows mean fluorescence intensity of four replicates. (b-c) Flow cytometry analysis on time course of conditioned EA.hy926 cells treated with S16-488 (b) and transferrin-633 (c) with or without 200 nM PA-SSSR. Treatment with PA-SSSR specifically reduces binding of S16-488 to endothelial cells but does not reduce the binding of transferrin-633, demonstrating that at least 60% of S16 cell binding occurs through the anthrax toxin receptors. MFI, mean fluorescence intensity. N=6 from two independent experiments. (d) Flow cytometry analysis on EA.hy926 WT and CMG2-/- cells treated with S16-488. Depletion of CMG2 significantly reduced S16 binding to S16 in a similar manner as CMG2 inhibition by PA-SSSR. (e) Conditioned EA.hy926 were incubated with 2 µM S16-488 and 10 ug/mL Tf-633 at 37C for 3 hours, then incubated in imaging buffer (SDM79 with 7.5 mM glucose and ProLong™ antifade) for 1 hr at 37C, then imaged by confocal microscopy. Representative images of two individual cell (top and bottom) are shown. Scale bar is 10 µM. Consistent colocalization between S16 and Tf in endosomes and in apparent perinuclear lysosomes was observed. These findings were consistent across n=20 cells from 2 independent experiments; in every cell, endosomal and lysosomal colocalization was also observed. ****, p < 0.0001; ns, not significant.

We then compared the ability of S16 to be internalized by Ea.hy926 cells in the presence and absence of 200 nM PA-SSSR. We chose PA-SSSR rather than WT because it has been demonstrated to be a better antagonist of angiogenesis than PA-WT.^*8*^ As shown in Figure 4b, S16-488 showed a time-dependent interaction with EA.hy926 cells that was drastically reduced (~ 60%) by co-incubation with PA-SSSR at concentrations high enough to compete with S16 for binding to cell-surface CMG2 ([PA] = 200 nM). In contrast, binding of Tf-633, was unaffected by the presence of PA-SSSR (Fig 4c). These data show that S16 and PA-SSSR share a cell-surface receptor. Since our *in vitro* FRET analysis indicates that S16 does not compete with PA for TEM8 binding (Supplementary Figure 6), TEM8 cannot be the major receptor responsible for S16 endocytosis.

To further validate that CMG2 mediates S16 endocytosis, we have developed a CMG2 knockout EA.hy926 cell line (CMG2-/-, Supplementary Figure 8) and evaluated its ability to internalize S16. CMG2-/- cells display significantly less S16-488 signal than WT EA.hy926 cells (50%; p<0.05; Figure 4d). This reduction is statistically indistinguishable from the reduction in S16-488 fluorescence observed in the presence of CMG2-saturating PA-SSSR concentrations (60%). Hence, we conclude that CMG2 mediates endocytosis of S16.

To gain insight into possible endocytic mechanisms involving CMG2, we used confocal microscopy to visualize the cellular uptake of S16-488 and Tf-633. We observed clear punctate intracellular colocalization of S16-488 and Tf-633 (Fig 4e, Supplementary Movie 1) in vesicles visible throughout the cell and concentrated around the nucleus in a pattern consistent with perinuclear lysosomes. Tf-633 is taken into cells via receptor-mediated clathrin-dependent endocytosis and can either be recycled through recycling endosomes or delivered to lysosomes. Hence, colocalization of S16 and transferrin in the same vesicular structures indicates that S16-488 is likely taken into cells via clathrin-dependent endocytic uptake, like transferrin, and therefore delivered to the same endosomes as transferrin. This mechanism of S16 endocytosis is consistent with the CMG2-mediated PA uptake, which also occurs via clathrin.*^16^*

## Conclusion

Together, the data presented in this work show that a small peptide-segment of the anti-angiogenic Col-IV fragment canstatin (α2 NC1) binds to CMG2 with sub-micromolar affinity, via the MIDAS (Fig 2), and profoundly inhibits endothelial cell migration. The near-complete inhibition of migration by S16 echoes the effect of other CMG2 antagonists*^8, 10, 13, 38^* and points to a critical role for CMG2 in this aspect of angiogenesis.

Our BLI, flow cytometry and confocal microscopy data show that CMG2 binds and mediates internalization of the peptide S16. Endothelial cells participate in ECM remodeling during angiogenesis, and are known to induce proteolytic cleavage of ECM materials and resulting release of ECM peptides. Such peptide cleavage products are then available for interaction with endothelial cell surface receptors, including CMG2. Indeed, many of these interactions and their subsequent signaling effects have been documented previously.^*40*^ While there is no evidence that S16 itself is a physiological ligand of CMG2, peptides containing relevant portions of the S16 primary amino acid sequence could, like other peptide proteolytic products, be available for binding and subsequent endocytosis by CMG2 during the process of ECM remodeling. A role for CMG2 in ECM homeostasis and uptake of ECM molecules and/or fragments has been previously proposed*^2, 15-17^*, particularly in the context of Col-VI.*^2^* Data presented here show that this CMG2-mediated process is not unique to Col-VI or Col-VI fragments, and may be a consistent cellular response to other ECM materials and/or peptides that also bind CMG2. CMG2-mediated endocytosis of a CMG2-binding molecule, as observed here, is consistent with the established role of CMG2 in mediating clathrin-dependent endocytosis of the anthrax toxin.*^2,41^* Indeed, co-localization of intracellular S16 with transferrin, as observed here, suggests that endocytosis of ECM-derived peptides like S16 could utilizes the same mechanism as endocytosis of anthrax toxin.*^41–44^*

Together, the findings included here suggest that CMG2 could bind and internalize Col-IV NC1 fragments, as is observed for S16, resulting in their eventual lysosomal degradation. This process provides one potential functional explanation for anti-angiogenic effects of CMG2 targeting. CMG2 interaction with, and endocytosis of, anti-angiogenic peptide fragments and/or protein domains such as arresten and canstatin would be expected to reduce their local concentrations, and thus, their anti-angiogenic impact. CMG2-binding molecules such as PA, S16, and small molecules*^10, 38^* may thus inhibit CMG2-mediated endocytosis of anti-angiogenic fragments and therefore magnify their anti-angiogenic impact. More work is needed to examine the mechanistic function of CMG2 in cell migration and to identify possible additional molecules involved in this process, which could be additional therapeutic targets for modulation of angiogenesis.

## Acknowledgement

The plasmid pGEX4T1-CMG2-GST was a generous gift from the John Collier lab.

