## Supplementary material for "A canstatin-derived peptide provides insight into the role of Capillary Morphogenesis Gene 2 in angiogenic regulation and matrix uptake": supplentary figures

**Supplementary Note 1:**

It is important to note that, consistently, the observed association of S16 to CMG2-loaded BLI sensors resulted in a negative binding signal, and the dissociation resulted in a positive signal. This is sometimes observed in BLI interaction data, and the built-in software feature to “flip” the data was used prior to generating the binding curve fits.

There are two likely reasons for the negative association, knowing that the interferometry signal originates as a function of sensor “thickness”: 1) S16 binding causes a large conformational rearrangement in CMG2 that results in a reduced hydrodynamic radius, and this “thickness reduction” owing to the conformational change outweighs the “thickness increase” from S16 binding (a small peptide); or 2) in our BLI assay, CMG2 is actually binding to S16 aggregates, likely on the order of 50 or more nanometers in diameter.

Interestingly, the observed association was inverted (negative association signal, and positive dissociation signal). While this does not impact the analysis, it is necessary to consider why this inverted binding signal might occur. In BLI, the readout is directly related to the change in effective size or thickness of the biosensor tip. Generally, as molecules bind to the sensor tip, the size of the sensor increases, causing a shift in the interference pattern of white light that yields a positive binding signal. However, if a small ligand (S16) binding CMG2 causes a large conformational change, which reduces the hydrodynamic radius of CMG2, it would be recognized as a negative signal. However, no direct evidence exists to support a binding-induced conformational rearrangement. An alternative evaluable solution is that large S16 aggregates are binding to CMG2. Indeed, both dynamic light scattering and fluorescence anisotropy suggest that S16 forms larger aggregates at concentrations > 10 uM, which is within the BLI assays range. Further, when we prepared S16 in a fashion that encourages monomerization, binding is not observed on BLI. However, this does not necessarily indicate that CMG2 binds only to S16 aggregates. Our BLI assay may not have sufficient sensitivity to detect the binding of the small S16 monomer (1600 Da). Regardless, the S16 peptide or aggregate binds CMG2 and elicits a profound phenotypic response in endothelial cells.

**Supplementary Figure 1. Membrane-based peptide array for CMG2 binding.** 15-mer peptides in a 10aa sliding window were synthesized directly coupled to a nitrocellulose membrane. This membrane was then probed with 250nM CMG2-biotin and read out using avidin-HRP. Left, low exposure imaging. Right, high exposure imaging. Proteins arrayed include: PA (blue), human Col-IV α1 (orange), human Col-IV α2 (green), and human fibronectin (black). Circled peptides include S16 (green) and U12 (red). Peptide in yellow circle is from triple-helical region of Col-IV α2, and thus was not further characterized here. Binding evident in PA-array is not surprising. Binding evident in fibronectin-array is currently being further validated and investigated.

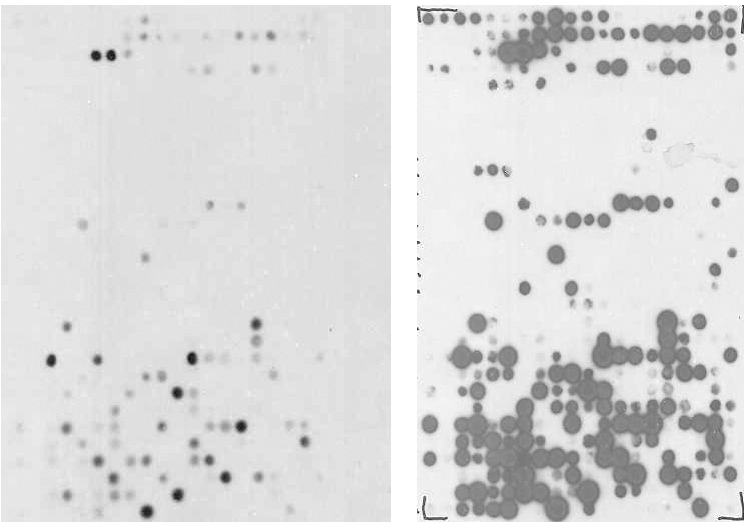

A

B

C

D

E

G

F

I

H

J

M

L

N

Q

P

R

S

O

K

T

U

V

W

X

G

Y

\

[

]

\#

1

2

3

4

5

6

7

8

9

1

0

1

2

3

4

5

6

7

8

9

2

0

A

B

C

D

E

G

F

I

H

J

M

L

N

Q

P

R

S

O

K

T

U

V

W

X

G

Y

\

[

]

\#

1

2

3

4

5

6

7

8

9

1

0

1

2

3

4

5

6

7

8

9

2

0

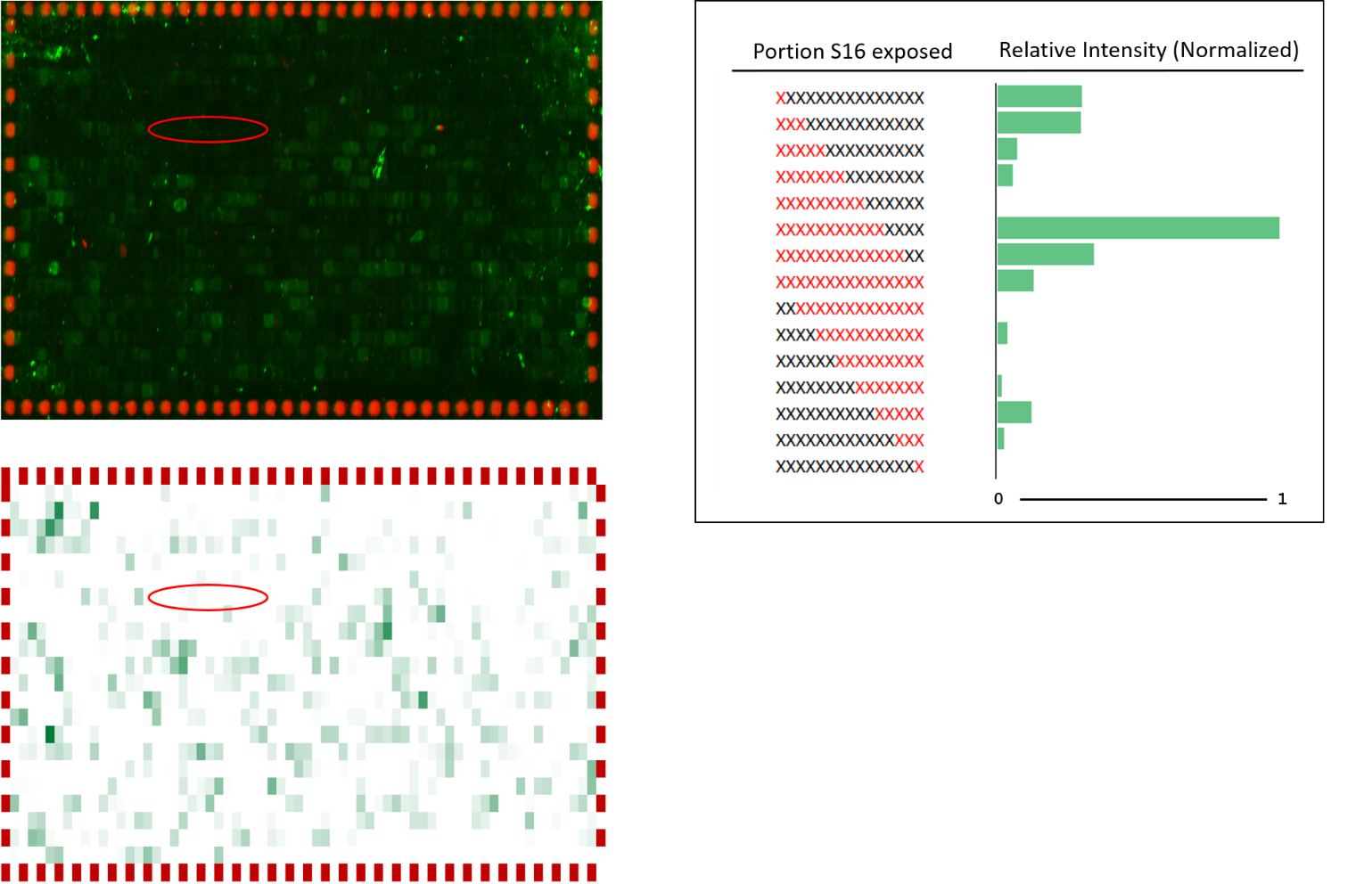

**Supplementary Figure 2. PEPperPRINT peptide array for CMG2 binding.** 15-mer peptides in a 2aa sliding window were synthesized with N- and C-terminal flanking linkers (GSGSGSG) in duplicate and coupled to a polyethylene-based graft copolymer. This array was then probed with 500 nM CMG2-GST-biotin and read out using an anti-GST-Dylight-800 conjugate. Top left, raw array image. Bottom left, heat map used to determine binding. Sequences including portions of S16 are circled in red. Proteins arrayed include: human Col-IV α1, human Col-IV α2, human laminin α1, α2 and α3, and selected epitopes from PA. Though S16-representing peptides did not exhibit high intensity compared to some other peptides assayed in this array, a clear binding pattern was observed in the S16 epitope (right, exposed amino acids represented in red). Binding evident in other proteins assayed is under current investigation.

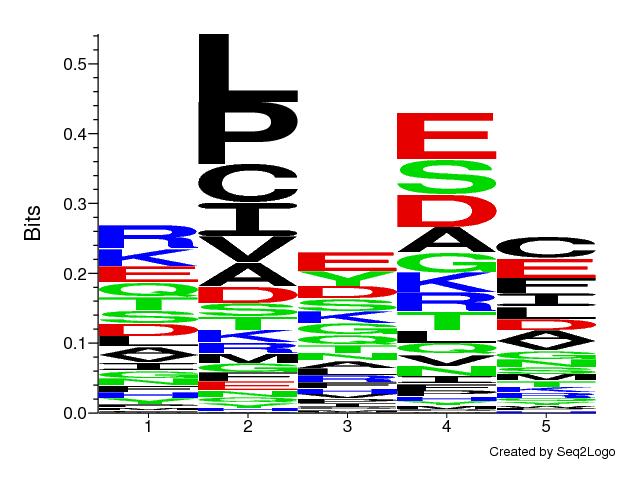

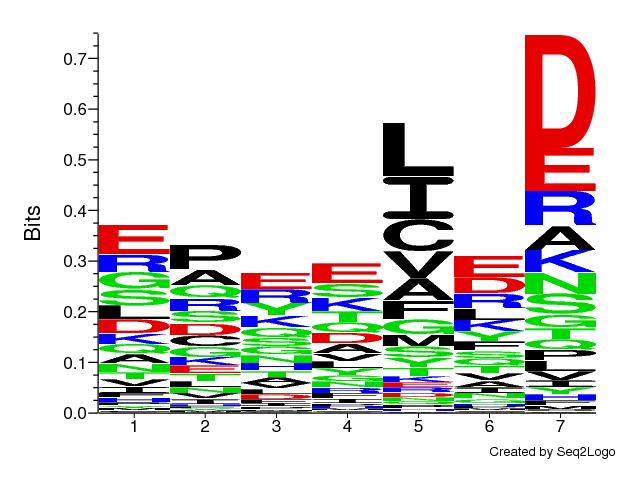

**Supplementary Figure 3. Identification of a potential CMG2-binding motif using PEPperPRINT array.** Top hits (65 peptides, false hits identified by visual inspection and removed) from array were pooled and subjected to clustering analysis with GibbsCluster-2.0. Top, 5-residue motif, with no insertions or deletions allowed. Bottom, 7-residue motif with up to 3 insertions and deletions allowed. Consistent motif consists of an enriched acidic position, preceded two positions by a hydrophobic/aliphatic residue.

U12

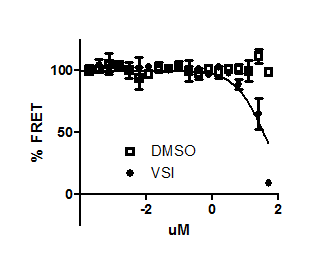

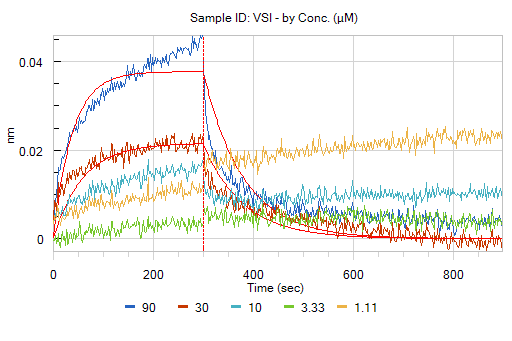

**Supplementary Figure 4. FRET and BLI demonstrate comparatively weak interaction of U12 with CMG2.** (top) PA-CMG2 competition FRET assay demonstrates an IC50 ≈ 35 µM for U12, over an order of magnitude weaker than S16. (bottom) Representative BLI sensorgram and fit for U12 interacting with CMG2-loaded SA sensors. Legend shows U12 concentration in µM. Signal/noise was too low for U12 concentrations <30 µM to be included in fit. Parameters for this fit are: K_d_ = 83 ± 6 µM, k_on_ = 150 ± 10 (Ms)^-1^, k_off_ = 0.0125 ± 0.0002 s^-1^. With low signal/noise fit quality is poor, but it is clear from replicates that K_d_ > 30 µM, in agreement with the FRET IC50.

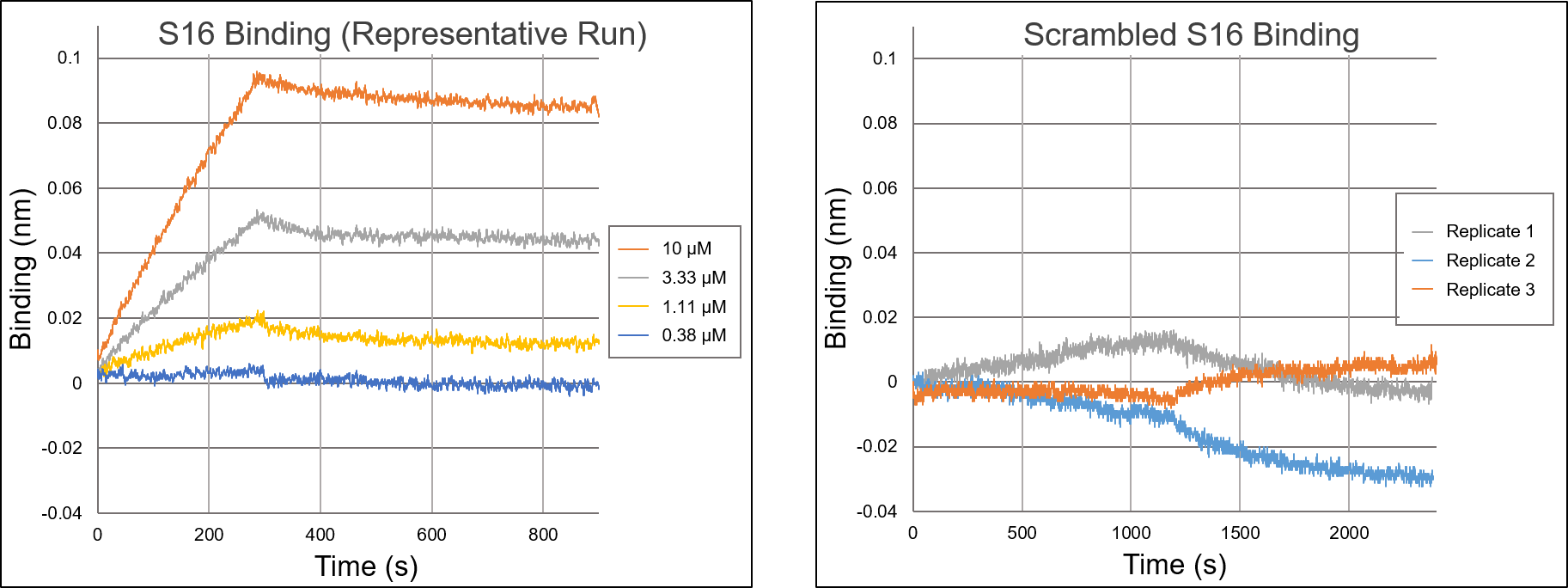

**Supplementary Figure 5. BLI analysis of CMG2**–**scrambled S16 interaction.**

A scrambled peptide, made by synthesizing a randomly shuffled S16 sequence, was tested on BLI for binding to CMG2. Assay conditions used were identical to those tested with S16–CMG2. Binding was compared across 3 independent experiments at 30 µM scrambled S16. In all experiments, the magnitude of the scrambled peptide signal was found to be minimal compared to that of S16, and data did not fit any binding models (shown above). Additionally, we note that scrambled S16 association and dissociation was assayed on a longer timescale (1200s) than the S16 assays. Any drift observed between samples is likely due to nonspecific, hydrophobic interactions of scrambled S16 with the sensor. These data provide evidence that the S16–CMG2 interaction is sequence-specific.

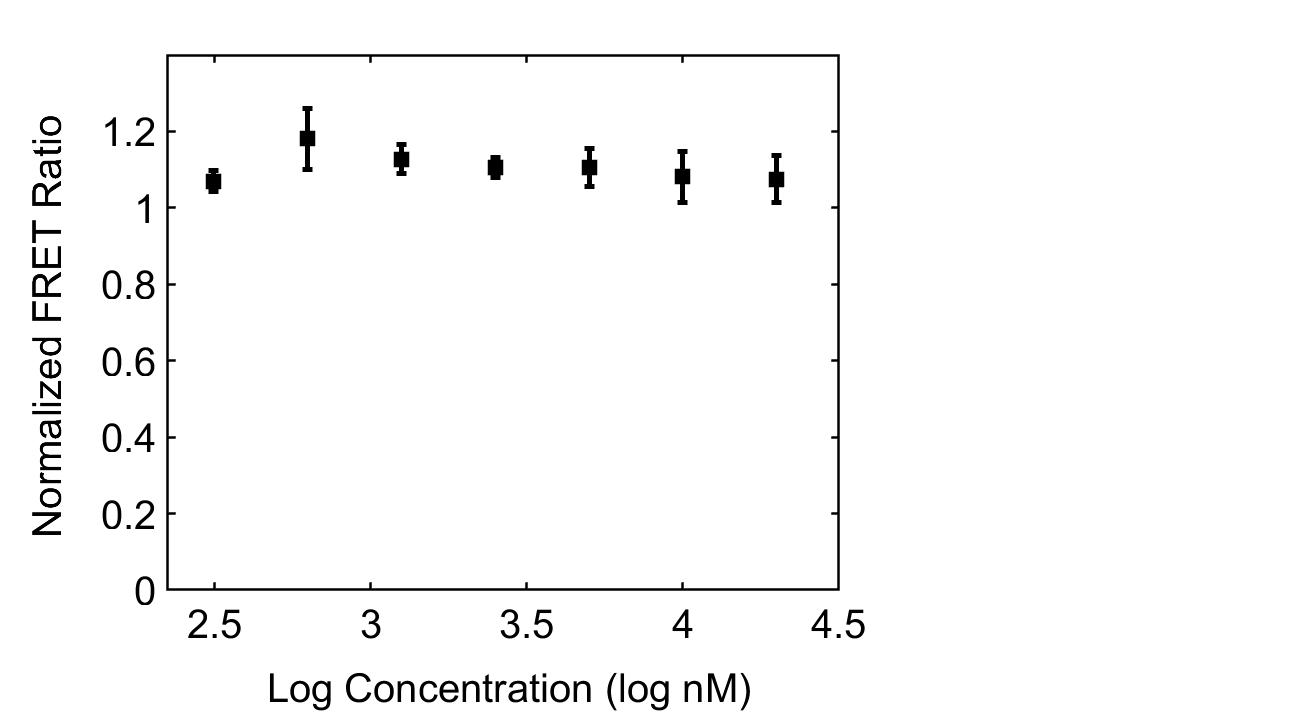

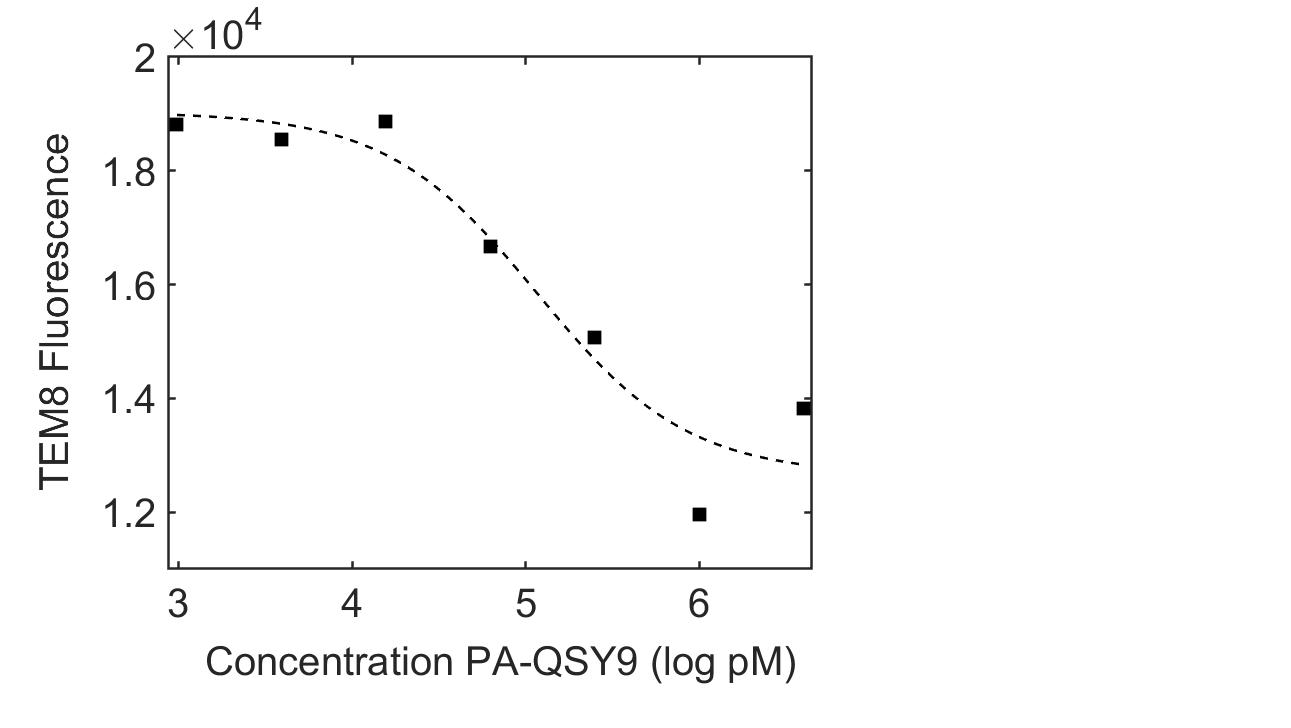

**Supplementary Figure 6. S16 does not interfere with TEM8-PA binding.** (Left) Inhibition of TEM8-GFP and PA-546 FRET was negligible at S16 concentrations between 313 nM and 40 μM, differing from that of CMG2–PA as shown in figure 2. TEM8-GFP and PA-546 mixed with DMSO vehicle only was used as normalization control. (Right) Activity of TEM8-GFP used in FRET inhibition assay was validated by measuring quenching by PA-QSY9. Measured K_D_ was approximately 100 nM (116 ± 111 nM) , matching published affinity for TEM8–PA.

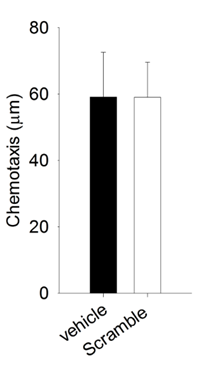

**Directional Migration (mm)**

**Supplementary Figure 7. EA.hy926 migration with scrambled S16 peptide.** The scrambled peptide is composed of the same amino acids as S16, but in a randomized sequence. Quantified directional migration of EA.hy926 cell migration in 0.3% DMSO (vehicle) and 10μM scrambled peptide had no significant difference. This data indicates that unlike S16, the scramble s16 has no effects on endothelial cell migration towards serum.

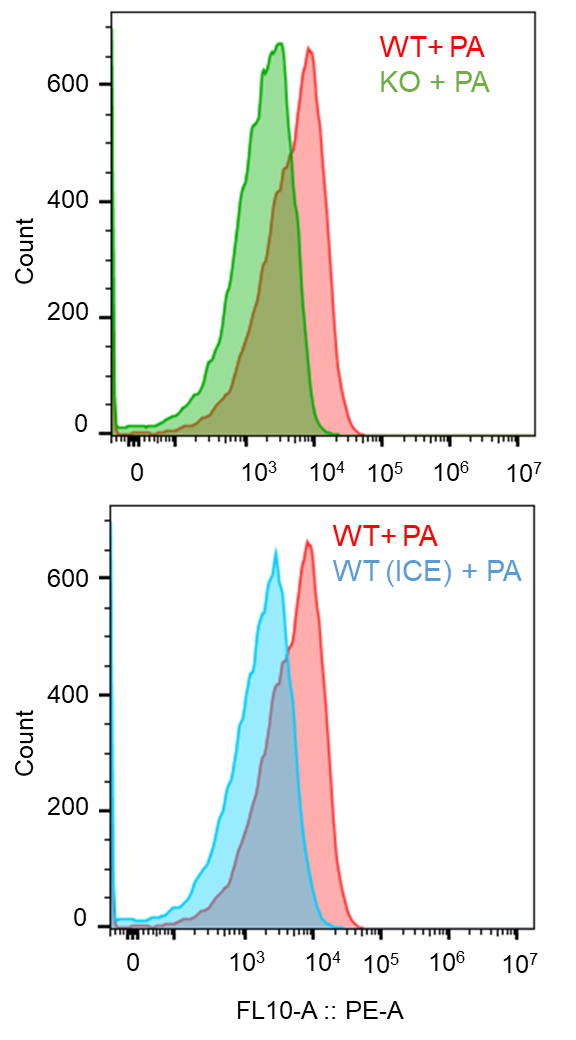

PA-Cy5 Intensity

**Supplementary Figure 8. EA.hy926 CMG2 -/- characterization via PA uptake assay.** CMG2 knockout in EA.hy926 cells was confirmed by PA uptake assays. EA.hy926 WT and CMG2 -/- cells were treated with 200pM of PA-WT-Cy5 conjugate, and the PA signal was measured via flow cytometry (Top). As observed above, CMG2-/- peak (green) is shifted to the left when compared to the WT peak (red). WT cells treated with PA on ice (blue) was also used as a control to compared with WT cells with PA at 37^o^C (Bottom). Endocytosis is expected to be stalled when cells are placed on ice. Data show that the peak position of CMG2-/- and WT cells on ice are very similar, indicating that CMG2-/- does not uptake PA at the concentration that only targets CMG2.

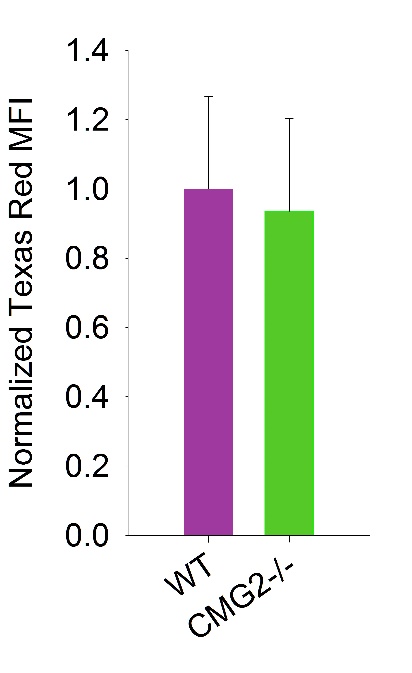

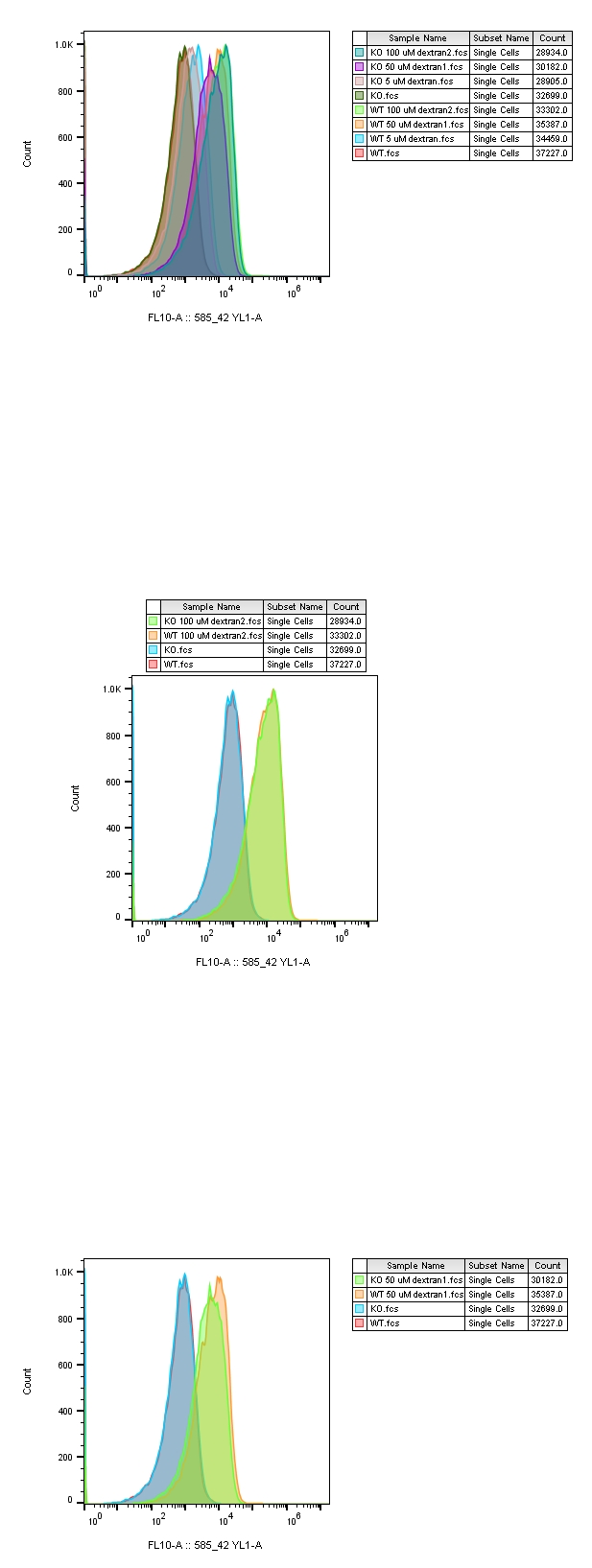

WT unstained

CMG2 -/- unstained

WT + Dextran

CMG2-/- + Dextran

**Supplementary Figure 9. Fluid phase uptake contributes to the non-specific uptake of S16.** Dextran uptake in both EA.hy926 WT and CMG2-/- cells. The two cell lines were pulsed with 100 µM Dextran Texas Red conjugate on ice for 1 hour, then chased at 37C^o^ with 5% CO_2_ for 4 hours. Texas Red signal was measured by flow cytometry. The data show that the peak shift in the Texas Red channel was similar between WT and CMG2-/- cells (Top left). Quantified data shows the mean fluorescent intensity difference between the two cell lines was insignificant (Top right). Therefore, CMG2 KO does not significantly affect rates of endocytosis in the cell and decreases in S16 uptake are via some other mechanism, most likely via abolished S16–CMG2 interactions. Additionally, these data show that non-specific uptake of material surrounding the cells is independent from CMG2-mediated endocytosis and can contribute to nonspecific S16 uptake.
